# Genetic basis of transgressive segregation in rice heading phenotypes

**DOI:** 10.1101/257766

**Authors:** Yohei Koide, Takashi Uchiyama, Yuya Ota, Shuntaro Sakaguchi, Ayumi Tezuka, Atsushi J. Nagano, Seiya Ishiguro, Itsuro Takamure, Yuji Kishima

**Affiliations:** Research Faculty of Agriculture, Hokkaido University, Kita-9 Nishi-9, Kita-ku, Sapporo, 060-8589, Japan; Faculty of Agriculture, Ryukoku University, 1-5 Yokotani, Seta Oe-cho, Otsu, Shiga 520-2194, Japan

**Keywords:** rice, transgressive segregation, extreme phenotype, days to heading, QTL

## Abstract

Transgressive segregation produces hybrid progeny phenotypes that exceed parental phenotypes. Unlike heterosis, extreme phenotypes caused by transgressive segregation are heritably stable. We examined transgressive phenotypes of flowering time in rice. Our previous study examined days to flowering (heading; DTH) in six F2 populations for which the parents had distal DTH, and found very few transgressive phenotypes. Here, we demonstrate that transgressive segregation in F2 populations occurred between parents with proximal DTH. DTH phenotypes of the A58 × Kitaake F2 progenies frequently exceeded those of both parents. Both A58 and Kitaake are *japonica* rice cultivars adapted to Hokkaido, Japan, which is a high-latitude region, and have short DTH. Among the four known loci required for short DTH, three loci had common alleles in A58 and Kitaake, and only the one locus had different alleles. This result indicates that there is a similar genetic basis for DTH between the two varieties. We identified five new quantitative trait loci (QTLs) associated with transgressive DTH phenotypes by genome-wide single nucleotide polymorphism (SNP) analysis. Each of these QTLs showed different degrees of additive effects on DTH, and two QTLs had epistatic effect on each other. These results demonstrated that genome-wide SNP analysis facilitated detection of genetic loci associated with the extreme phenotypes and revealed that the transgressive phenotypes were produced by exchanging complementary alleles of a few minor QTLs in the similar parental genotypes.

## INTRODUCTION

The range of phenotypic variation in a quantitative trait depends on its genetic complexity (Alonso-Blanco and Mendez-Vigo 2014; Huang and Han 2014). Cross hybridizations often produce progenies with wider phenotypic variation than their parents, which is referred to as transgressive segregation (Rick and Smith 1953; Harlan 1976; de Vicente andTanksley 1993). Unlike heterosis, the extreme phenotypes that occur as a result of transgressive segregation can be fixed after the second filial generation (F2). Such extreme phenotypes can have important roles in evolution (Rieseberg *et al.* 2002; Dittrich-Reed and Fitzpatrick 2013). From a breeding perspective, this phenomenon has also strongly contributed to crop and animal improvements (Vega and Frey 1980; Tanksley and McCouch 1997). However, little is known about the genetic basis of transgressive segregations, which are associated with phenotypic improvement of useful traits in crops.

Days to heading (DTH) determines the regional adaptability of rice (*Oryza sativa* L.), which is cultivated widely in tropical and temperate regions (Hori *et al.* 2016). DTH is an important agronomic trait that controls flowering time in rice. Flowering time is a complicated trait in many crops, and the genetic basis of DTH has been well studied in rice; to date, 14 quantitative trait loci (QTLs) were identified based on natural variation and isolated by map-based cloning strategies (Ebana *et al.* 2011; Hori *et al.* 2016; Brambilla *et al.* 2017). We previously examined DTH in rice using six different F2 populations derived from crosses between Kokusyokuto-2 (a Hokkaido landrace denoted as A58) with a short DTH (81 days) used as the seed parent and six varieties with long DTH (114-126 days) used as the pollen parents (Ota *et al.* 2014). Most F2 plants from all six crosses showed intermediate DTH that fell within the parental ranges. Ota *et al.* (2014) found some plants with shorter DTH than A58 from the A58 × Kasalath F2 population; only this F2 population had some individuals with shorter DTH relative to those of the parents, and the other five F2 populations did not exhibit such extreme phenotypes. In the plants with shorter DTH, we identified a genetic interaction *(Ghd7* from A58 and *Ehd1* from Kasalath) that contributed to the extreme phenotypes produced by the cross of the distantly related parents (Ota *et al.* 2014).

Here, we were interested in determining how the range of phenotypic variation is produced and whether extreme phenotypes can be produced when both parents used in the cross have proximal phenotypes. The progenies from parents with the same genotypes would have very narrow phenotypic variation, while certain range of the phenotypic variation like transgressive phenotypes should be expected in the F2 generation from parents with different genotypes that could coincidentally cause similar phenotypes. By testing these predictions, we may be able to identify the unknown genetic entities that produce extreme phenotypes.

We specifically focused on phenotypic variations in DTH of a population derived from a cross between two closely related varieties, A58 and Kitaake (an improved variety), both of which are adapted to Hokkaido, a northernmost rice cultivation area. Compared with the progenies of distantly related parents with distal DTH phenotypes, more A58 × Kitaake progenies had extreme short or long DTH phenotypes relative to the parents. We evaluated the genetic causes of transgressive segregations of both early and late DTH observed in this segregating population. First, known genes associated with short DTH were evaluated in A58, Kitaake, and their progenies to determine if transgressive phenotypes were produced. Subsequently, we performed genome-wide single nucleotide polymorphism (SNP) analysis to detect unknown QTLs associated with extremely short or long DTH. The results obtained here demonstrated that a relatively small number of minor QTLs and their epistatic interactions produced transgressive segregation in DTH. Important genetic properties of the extreme heading phenotypes caused by transgressive segregation are discussed.

## MATERIALS AND METHODS

### Genetic stocks

A rice landrace from Hokkaido, A58, and an improved variety of Hokkaido, Kitaake, were used as parents. A58 seeds were obtained from seed stocks at the Plant Breeding Laboratory in Hokkaido University. Kitaake seeds were obtained from the genebank at the Agricultural Research Department of Hokkaido Research Organization. A58 was crossed with Kitaake to obtain F1 seeds. A total of 248 F2 plants were obtained from self-pollination of the F1 plants. From the 248 F2 plants, 132 were randomly selected to obtain F3 populations. These F3 populations were used for genetic analysis of DTH using DNA markers in the *Hd1* locus, which is a major locus that affects DTH in rice (Yano *et al.* 2000). Of the 132 F3 populations, 15 individuals that showed early DTH and had a fixed *Hd1* genotype were selected as early-heading populations. Similarly, 15 individuals that showed late DTH and had a fixed *Hd1* genotype were selected as late-heading populations. Plants in these two selected populations were self-pollinated to produce F4 lines. The genotypes of the 15 early and 15 late lines in the F4 generation were determined by genome-wide SNP analysis (Figure S1).

### DTH analysis

Plants were grown in an experimental paddy field at Hokkaido University, Sapporo, Japan (43.1 N). For F2 and F3 populations, DTH was measured in 2013 and 2014, respectively. DTH for the F4 generation was measured in 2015 and 2016 as the number of days from sowing to emergence of the first panicle of a plant. Average DTH of the F3 and F4 populations were calculated from the values of five or six plants.

### Genotyping and sequencing

Genomic DNA was extracted from leaf samples using Plant DNAzol (Invitrogen, Carlsbad, CA, USA). To genotype the *Hd1* locus, two primers, Hd1L (5′-CGA CGT GCA GGT GTA CTC CG-3′) and Hd1R (5′-AAT CTG TGT AAG CAC TG ACG-3′), were used based on the *Hd1* sequence. Genome-wide SNPs were detected by double digest restriction site-associated DNA sequencing (ddRAD-Seq) (Baird *et al.* 2008; Peterson *et al.* 2012), which began with DNA library preparation using the restriction enzymes *BglII* and *EcoRI.* Sequencing was performed with 51 bp single-end reads in one lane of a HiSeq2000 Sequencer (Illumina, San Diego, CA, USA) by Macrogen (Seoul, South Korea). The ddRAD-sequencing reads were trimmed with Trimmomatic ver 0.33 (Bolger *et al.* 2014) with the following parameters: LEADING:19 TRAILING:19 SLIDINGWINDOW:30:20 AVGQUAL:20 MINLEN:51. The trimmed reads were mapped to a RAD reference for the Os-Nipponbare-Reference-IRGSP-1.0 using Bowtie 2 (Langmead and Salzberg 2012) with a default parameter setting. To build RAD loci, we used the ref_map.pl pipeline in Stacks ver. 1.29 (Catchen *et al.* 2011). All RAD-Seq procedures were carried out by Clockmics, Inc. (Izumi, Osaka, Japan). A total of 1,402 SNPs between parental varieties were detected by ddRAD-Seq; among these SNPs, 634 were considered reliable after filtering SNPs that appeared in more than 80% of F4 plants.

PCR amplicons for the four previously identified genes involved in DTH *(Hd1*, Hd2/OsPRR37, *E1/Hd4/Ghd7*, and *Hd5/DTH8)* were purified using a NucleoSpin Gel and PCR Clean-up kit (Macherey-Nagel, Düren, Germany). Purified samples were sequenced in both directions using a Big Dye Terminator Cycle Sequencing kit (Applied Biosystems, Foster City, CA, USA) on an ABI310 automatic sequencer (Applied Biosystems). Sequence alignment was performed using CLUSTAL W 2.1 (Thompson *et al.* 1994). The primers used for sequencing the four genes were as follows: *Se1/Hd1* [5′-CGA CGT GCA GGT GTA CTC CG-3′ and 5′-AAT CTG TGT AAG CAC TG ACG-3′], *Hd2/OsPRR37* [5’-TCT TTC TGA TGG CTG TCT GC-3′ and 5′-GCC ATC GCG TAG GTA GGT AG-3′], *E1/Hd4/Ghd* [5′-GCT GGC TGG ACT TCA CTA CC-3′ and 5′-CAT GGG CCA CTT CTA AGA TCA-3′], and *Hd5/DTH8* [5′-CGG AGT TCA TCA GCT TCG TT-3′ and 5′-TGA CCA TGG TGT GAG TGT GA-3′].

### Transgressive index

We defined the transgressive index, which indicates the proportion of phenotypic differences between both parents and the phenotypic range in the F2 population. The transgressive index was calculated by dividing the width of the distribution of DTH in the F2 population by the parental DTH difference.

### Marker genotype value

Allelic effects of each of the six loci that influenced DTH of the A58 × Kitaake hybrid progenies were evaluated as marker genotype values (Goddard and Hayes 2007). Average DTH for each allele was calculated based on DTH data collected from all homozygous alleles in the 30 F4 population lines from 2016. Then, the central value was determined based on the two phenotypic averages obtained from each of both alleles at the same locus. The marker genotype value was equivalent to the difference between the central value and either allelic average DTH in the same locus.

### Data availability

All genetic stocks and sequence data are available on request. Sequence data for this study were deposited in DDBJ (DDBJ accession number XXXXX).

## RESULTS

### Variation and transgressive phenotypes of DTH in A58 × Kitaake progenies

Both A58 and Kitaake are adapted to the high-latitude area between 41.2 N and 45.4 N in Hokkaido, Japan, and consequently have photoperiod-insensitive, short DTH and are cold-resistant (Ishiguro *et al.* 2014; Ota *et al.* 2014). There was no significant DTH difference between these two varieties (t-test: A58, 81.2 ± 0.38; Kitaake, 80.5 ± 0.66; P = 0.19) (Figure 1A). DTH of these F2 plants were widely distributed from 69 to 87 days (Figure 1A), and the earliest plant DTH was equivalent to that of an extreme early-heading variety (Figure 1A).

**Figure 1.**
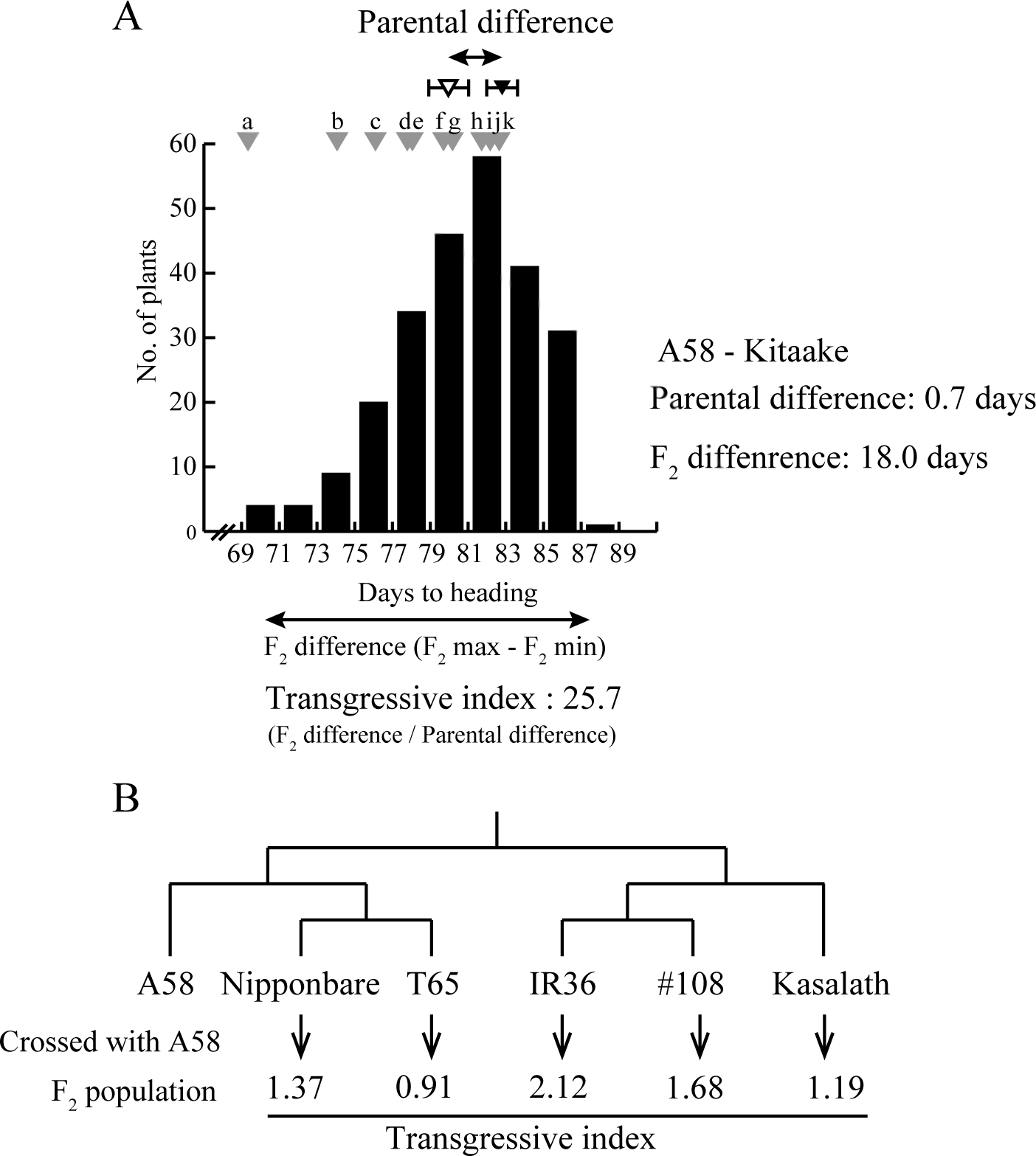
Transgressive segregation and genetic relationships between parental varieties. (A) Frequency distribution of DTH from A58 × Kitaake F2 plants. The transgressive index represents the ratio of the of the F2 population DTH distribution to the parental difference. The DTH difference between A58 and Kitaake was 0.7 days, and the DTH range in the F2 population was 18.0 days, which resulted in a transgressive index of 25.7. Standard DTH values of 11 varieties in Hokkaido are indicated by gray arrowheads: (a) Kitaibuki, (b) Hakucho-mochi, (c) Daichinohoshi (d) Hatsushizuku, (e) Hoshinoyume, (f) Kuiku180, (g) Hokuiku-mochi, (h) Nanatsuboshi, (i) Kirara397, (j) Hoshimaru, and (k) Gimpu. (B) Transgressive indexes of crosses between A58 and each of five other varieties. The phylogenetic relationships and the associated dendrogram for the five O. sativa varieties, Nipponbare (japonica), T65 (japonica), IR36 (indica), #108 (indica), and Kasalath (indica, Aus), are presented based on information provided in Takata et al. (2005). To calculate the transgressive index, DTH of parental varieties and F2 plants were calculated based on data from Ota et al. (2014).

The transgressive index of the A58 × Kitaake progenies was 25.7 (Figure 1A). In the six crosses between A58 and the other varieties that were more genetically distant from A58 than Kitaake (Ota *et al.* 2014), the transgressive indexes ranged from 0.91 to 2.12 (Figure 1B). The transgressive index of DTH indicated that the DTH of A58 × Kitaake F2 plants exceeded DTH of either parent, but such a strong transgressive segregation was not observed in the previously published crosses.

The DTH distribution in the A58 × Kitaake-derived F3 populations was similar to that of the F2 population (Figure S2). For further analysis, we selected 15 early- and late-heading F3 plants and developed two F4 populations (early and late) by self-pollination. Average DTH of early- and late-heading F4 populations were 63.8 ± 1.32 and 74.6 ± 0.99, respectively, in 2015, and 72.2 ± 1.32 and 80.0 ± 1.00, respectively, in 2016 (Figure 2). These differences between DTH of the early and late populations were significant (t-test, P < 0.001) throughout 2015 and 2016, and it was predicted that these two distinct populations were generated by new genetic interactions derived from the A58 × Kitaake cross.

**Figure 2.**
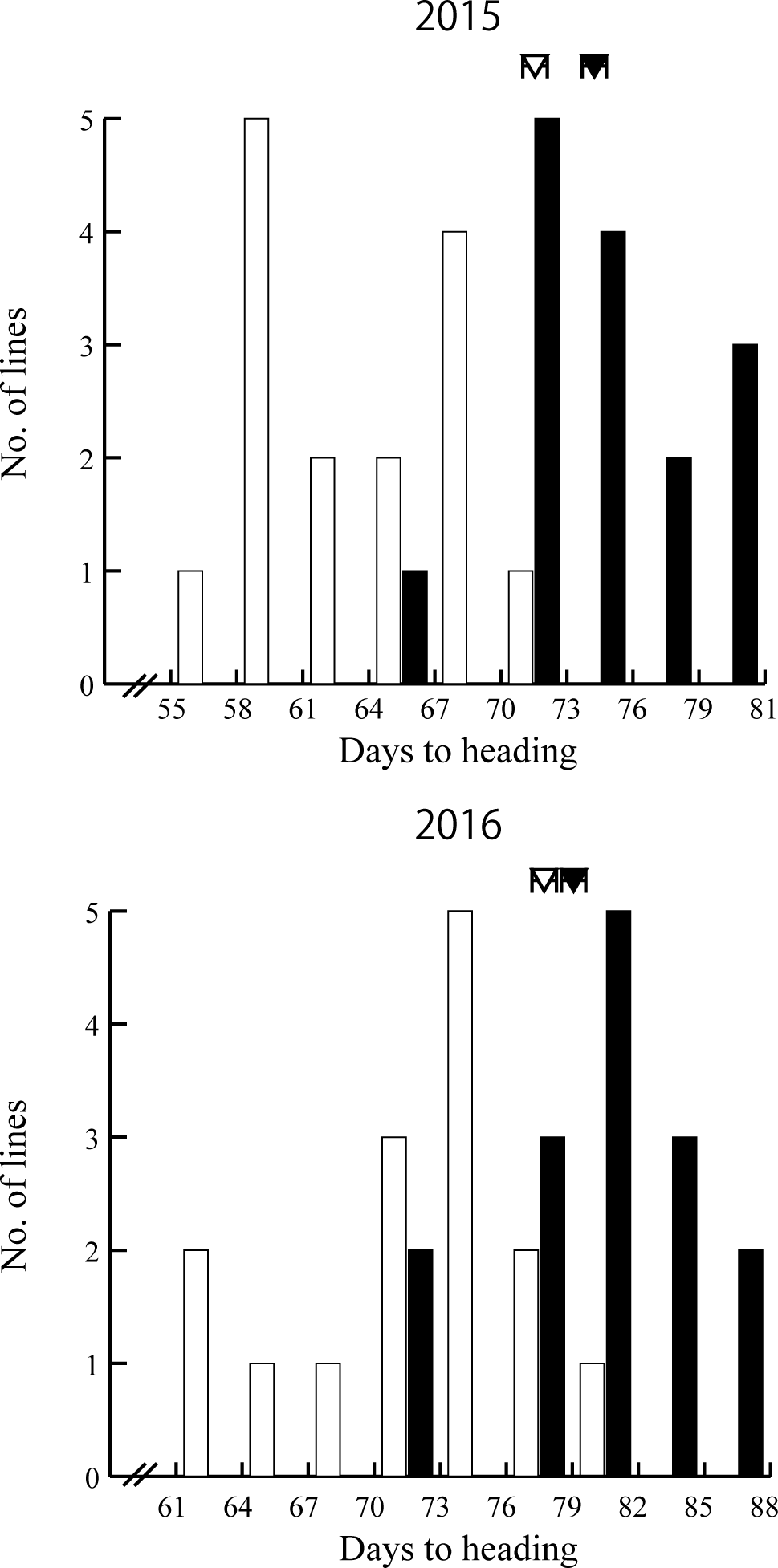
Frequency distribution of DTH in early- and late-heading F4 lines derived from the A58 Kitaake cross in 2015 and 2016. DTH of the 15 early- and 15 late-heading F4 lines selected in the F3 population was examined in the two years, 2015 and 2016. Early- and late-heading lines are indicated by white and black, respectively. Kitaake and A58 DTH are indicated by white and black arrowheads, respectively, with bars indicating S. E.

### Sequence analysis of genes that control DTH, and the effect of *Hd1* on DTH

Four loci (*E1/Hd4/Ghd7, Hd2/OsPRR37, Se1/Hd1*, and *Hd5/DTH8*) control DTH in varieties from Hokkaido, and their specific alleles facilitated adaptation by producing photoperiod-insensitive varieties with short DTH (Ichitani *et al.* 1997; Fujino and Sekiguchi 2005a; Fujino and Sekiguchi 2005b; Nonoue *et al.* 2008; Fujino *et al.* 2013; Koo *et al.* 2013). To confirm whether these four loci are related to the DTH differences observed in the A58 × Kitaake F2 population, we compared nucleotide sequences of these loci (Figure 3). Sequence analysis of *Hd1* showed the presence of polymorphisms, including a 312-bp insertion/deletion in A58 and Kitaake. This polymorphism in *Hd1* might have produced the DTH differences observed in the F2 population. In contrast to *Hd1*, no polymorphisms were detected in the other three loci *(E1/Hd4/Ghd7, Hd2/OsPRR37*, and *Hd5/DTH8)* in A58 and Kitaake (Figure 3).

**Figure 3.**
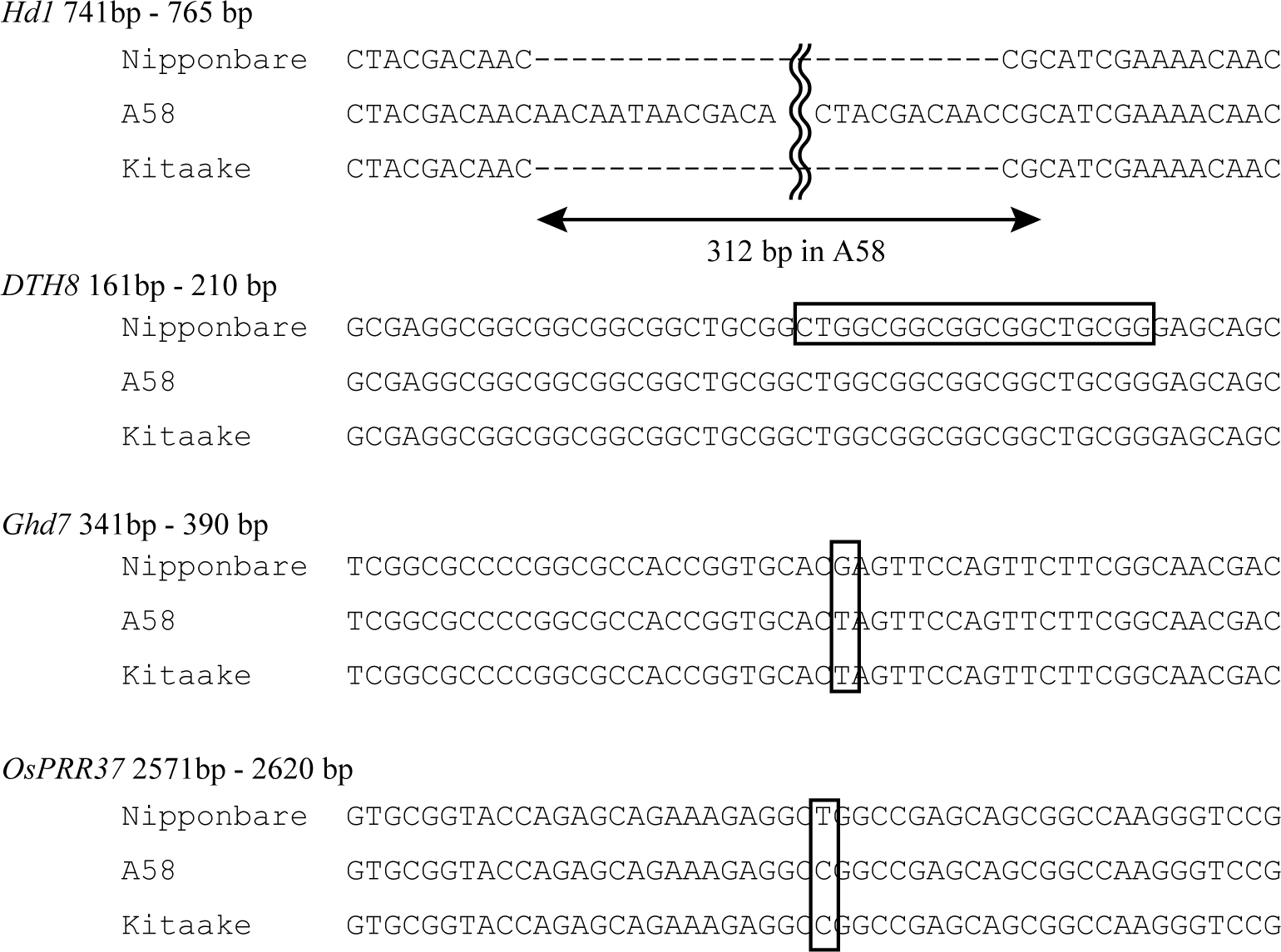
Comparisons of partial nucleotide sequences from Nipponbare, A58, and Kitaake for the four major loci that affect DTH in Hokkaido. The sequenced positions (based on Nipponbare) were selected using known polymorphisms among varieties in Hokkaido that were observed in previous studies (Ichitani et al. 1997; Fujino and Sekiguchi 2005a; Fujino and Sekiguchi 2005b; Nonoue et al. 2008; Fujino et al. 2013; Koo et al. 2013). Hd1 showed multiple differences between A58 and Kitaake; Kitaake possesses a functional allele that is also found in Nipponbare. In DTH8, a 19-bp segment (indicated by a rectangle) was deleted in most of the other Hokkaido varieties, but not in Nipponbare, A58, and Kitaake, from which we could not detect any polymorphisms. For Ghd7 and OsPRR37, SNPs observed in Nipponbare and the other two varieties are indicated by boxes.

In terms of the effect of the *Hd1* locus on the A58 × Kitaake F2 population, the average DTH in A58-type homozygous, heterozygous, and Kitaake-homozygous populations were 81.3 ± 0.36, 79.5 ± 0.38, and 78.8 ± 0.46, respectively (Table 1). The results showed that *Hd1* had a significant but small effect on DTH in this population (P < 0.001), which revealed that the extremely early phenotype of progenies was not fully explainable by only *Hd1*, and another factor(s) may be involved.

**Table 1.** The effect of Hd1 locus on days to heading in F2 population derived from A58 x Kitaake cross

| Marker |  |  | No. of plants |  |  | Average DTH |  |  | *P* |
| --- | --- | --- | --- | --- | --- | --- | --- | --- | --- |
| name | Chr. | Position | A58-type | Heterozygous | Kitaake-type | A58-type | Heterozygous | Kitaake-type |  |
| Hd1 | 6 | | 56 | 103 | 73 | 81.3 | 79.5 | 78.8 | $4 \times 10^{-4}$ |

### Detection of SNPs associated with extreme DTH phenotypes

If a QTL for DTH was located near an SNP, the SNP alleles tended to be associated with early- or late-heading populations. Genome-wide SNP analysis using ddRAD-Seq provided us a total of 634 reliable SNPs for 15 early and 15 late lines in the F4 populations (Figure 4). Among the 634 SNPs, 27 were detected as loci where the frequency of either the A58- or Kitaake-type allele was distorted in early- or late-heading populations (Table 2). Of these possible DTH phenotype-related SNPs, we focused on 19 SNPs that belonged to five clusters on Chs 1, 2, 4, 6, and 10 (Table 2 and Figure 4); these SNP clusters represented the chromosomal regions where QTLs for DTH were present.

**Figure 4.**
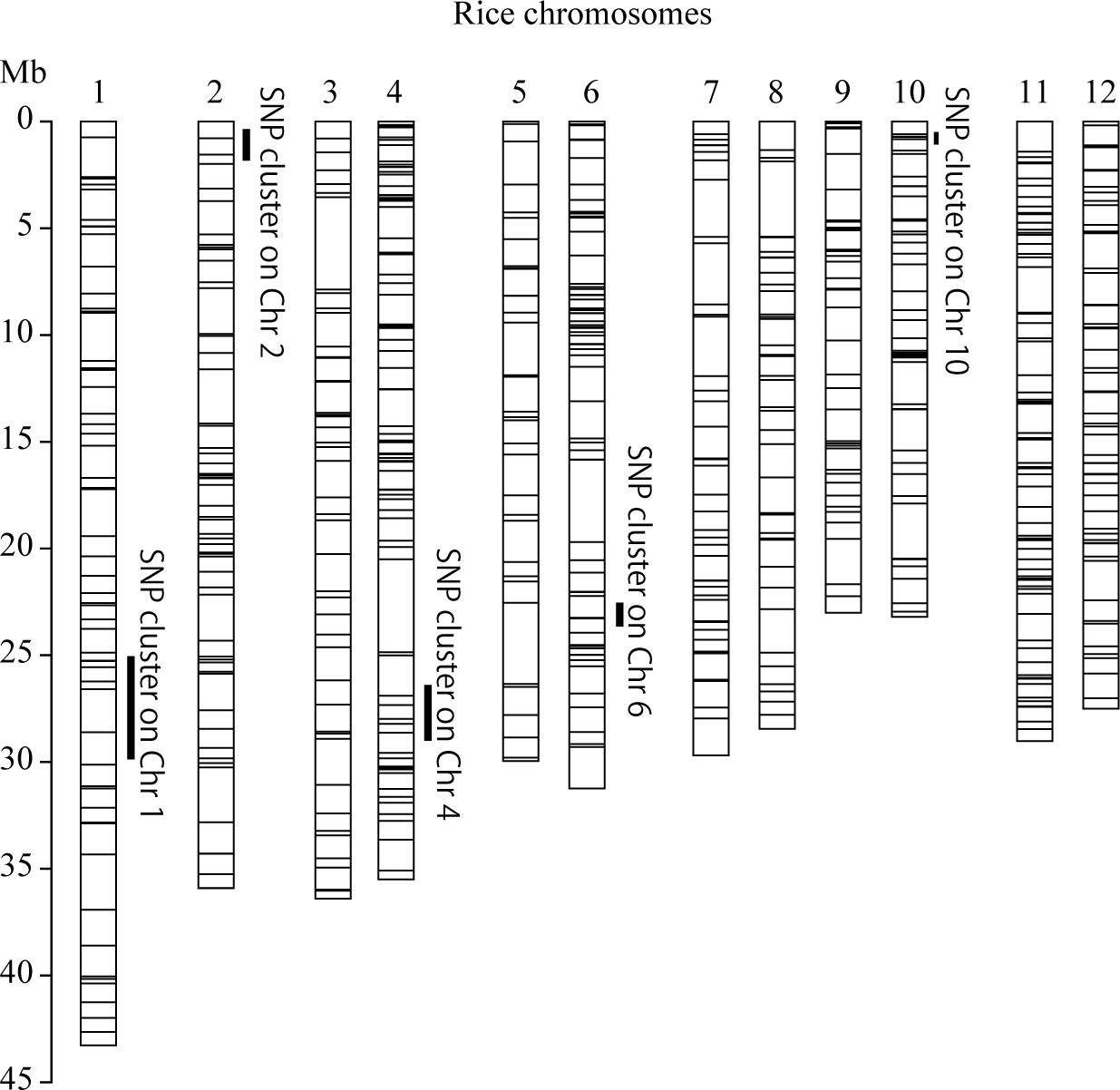
Chromosomal positions of five SNP clusters. Physical map positions of each SNP detected by ddRAD-Seq are shown by horizontal bars in each chromosome. Positions of SNP clusters that showed significant differences in allele frequency between early- and late-heading populations are indicated by vertical bars on the right side of each chromosome.

**Table 2.**
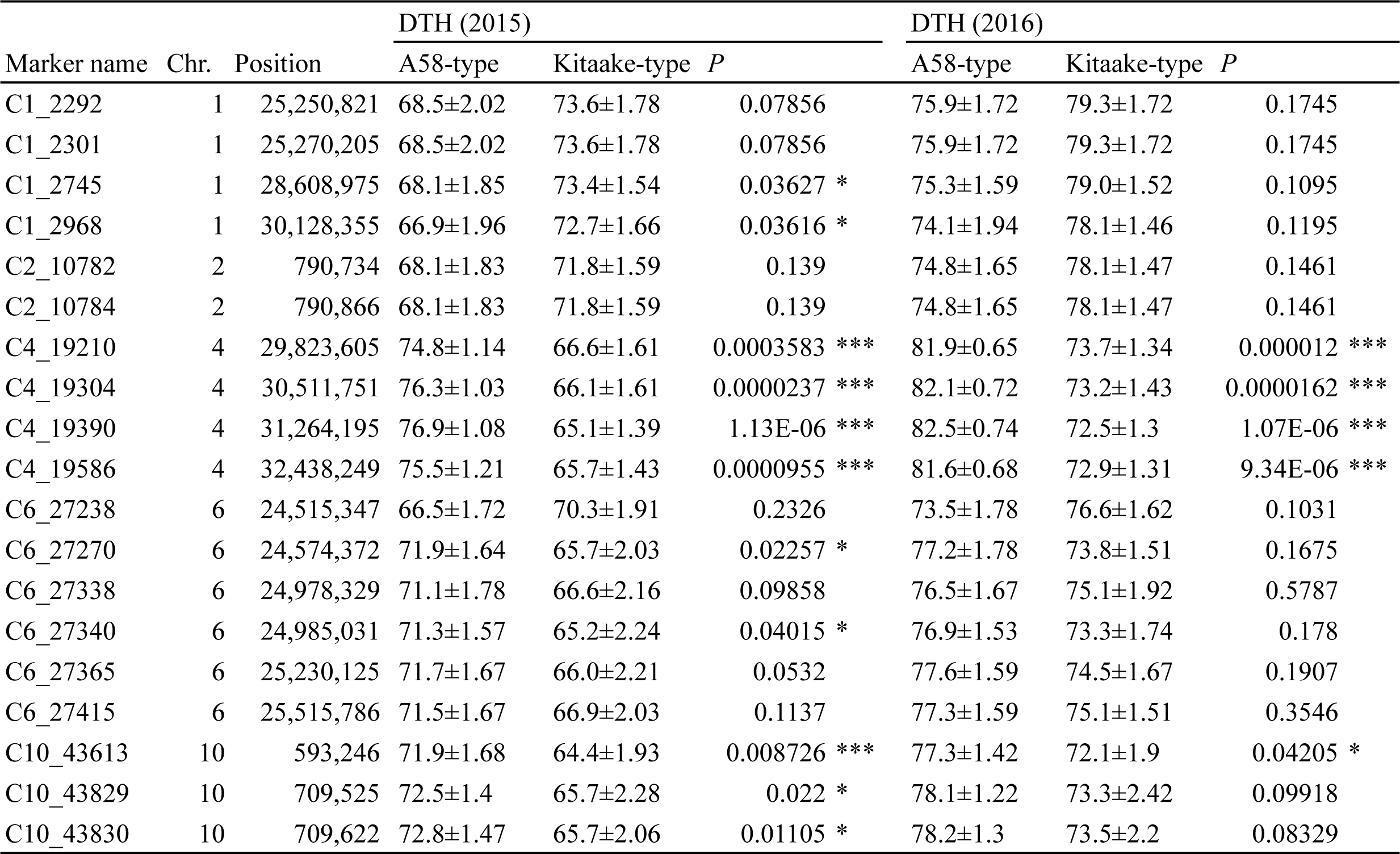
The effect of SNP clusters on days to heading in F4 lines derived from the cross between A58 and Kitaake

### Validation of relationships between SNP genotypes and DTH

Table 2 shows average DTH between two alleles of each of the five SNP clusters in the F4 lines examined in 2015 and 2016. Significant differences (P < 0.001) in DTH between A58- and Kitaake-type homozygous alleles were observed in SNPs on Ch 4 in both 2015 and 2016 (Table 2 and Figure S3). In the SNP cluster on Ch 10, significant differences in DTH between the two alleles were also observed, although the difference was small in 2016. SNPs on Chs 1 and 6 weakly significantly differed between the two alleles that were only observed in 2015. Among the five clusters, the weakest effect was detected in the Ch 2 cluster, which was not significant (P = 0.14), but still clearly discriminated the two alleles by 3 to 4 days. Overall, the order of the five SNP clusters based on additive effects on DTH was: Ch 4 > *(Hd1)* > 10 > 6 > 1 > 2. The Kitaake-derived alleles in the SNPs on Chs 4, 10, and 6 produced shorter DTH than the A58-derived alleles; alternatively, the A58-derived alleles on Chs 1 and 2 produced shorter DTH than the Kitaake-derived alleles.

Among the selected chromosomal regions (Chs 1, 4, 6, and 10), genetic interactions were tested using the F2 population (Figure 1). Among several combinations of possible epistatic interactions, a strong genetic interaction was identified in the SNPs on Chs 1 and 10 (Figures 5 and S4). The A58-derived alleles in the SNPs on Ch1 decreased DTH when they were combined with Kitaake-derived alleles in SNPs on Ch 10, but increased DTH when combined with A58-derived alleles in SNPs on Ch 10 (Figure 5). No known genes associated with DTH are located around these two chromosomal regions. These findings revealed that unknown genes from A58 and Kitaake caused epistatic interactions responsible for the transgressive early phenotype.

**Figure 5.**
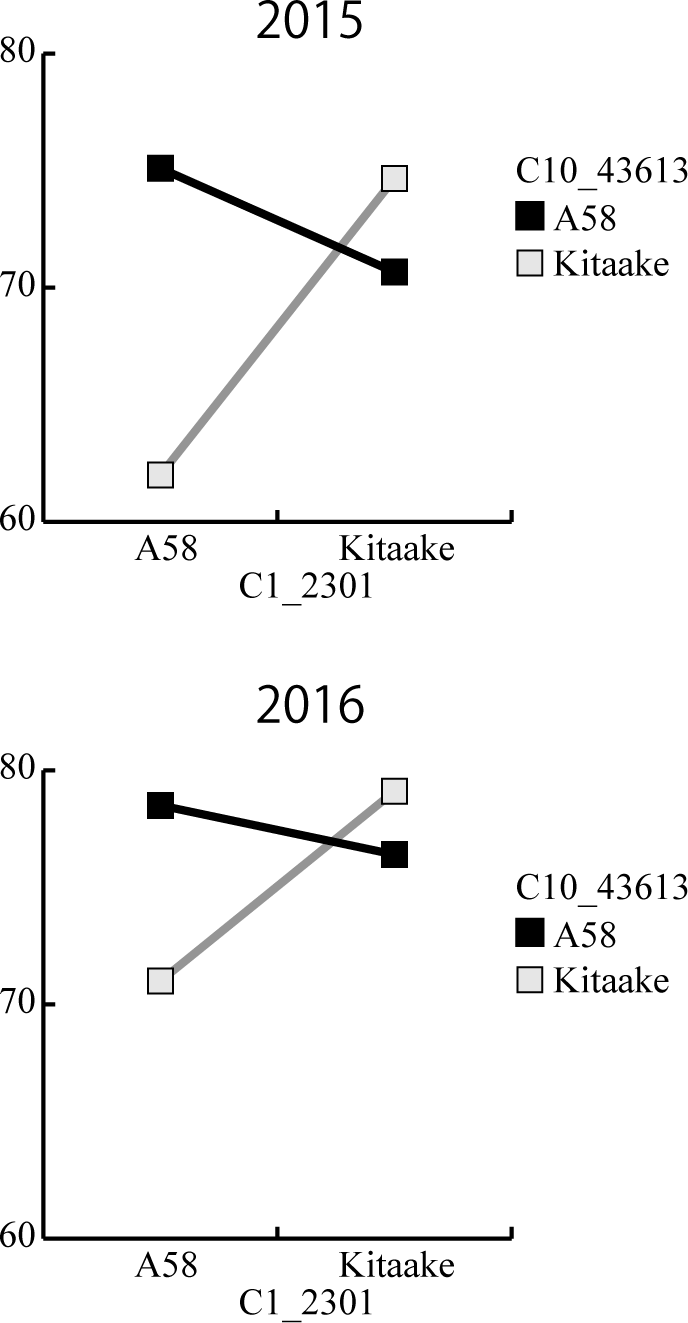
Epistatic interaction between SNPs on chromosomes 1 and 10 on DTH observed in 2015 and 2016. Average DTH values for the four combinations of genotypes with central SNPs (C1_2301 and C10_43613) in the clusters on Chs 1 and 10, which are indicated by squares. The case of Chs 1 and 10 were selected from all the combinations with Chs 1, 4, 6 and 10 (Figure S5). When the A58 SNP on Ch 10 (black line) and Kitaake SNP on Ch 10 (gray line) were respectively coupled with the different parental SNPs, epistatic (allelic) interactions occurred; in particular, the combination of the A58 allele on Ch 1 and Kitaake allele on Ch 10 resulted in the shortest DTH.

Loci weighted by marker genotype values based on DTH data from 2016 are shown in Figure 6. Among a total of 30 F4 lines, nine harbored homozygous alleles at all six loci (the five QTLs and *Hd1).* These nine lines were sorted by DTH (Figure 6). Based on marker genotype values, direction of allelic effect, and the numbers of the alleles with an effect, the order based on DTH was reasonable, although it was not identical to the expectation, because of possible genetic interactions or unknown genes. The short DTH lines tended to have more alleles with a short DTH effect, whereas the long DTH lines contained more alleles with the opposite effect. Therefore, the extreme phenotypes produced by transgressive segregation might be defined by allelic composition with different phenotypic effects occurring in either direction.

**Figure 6.**
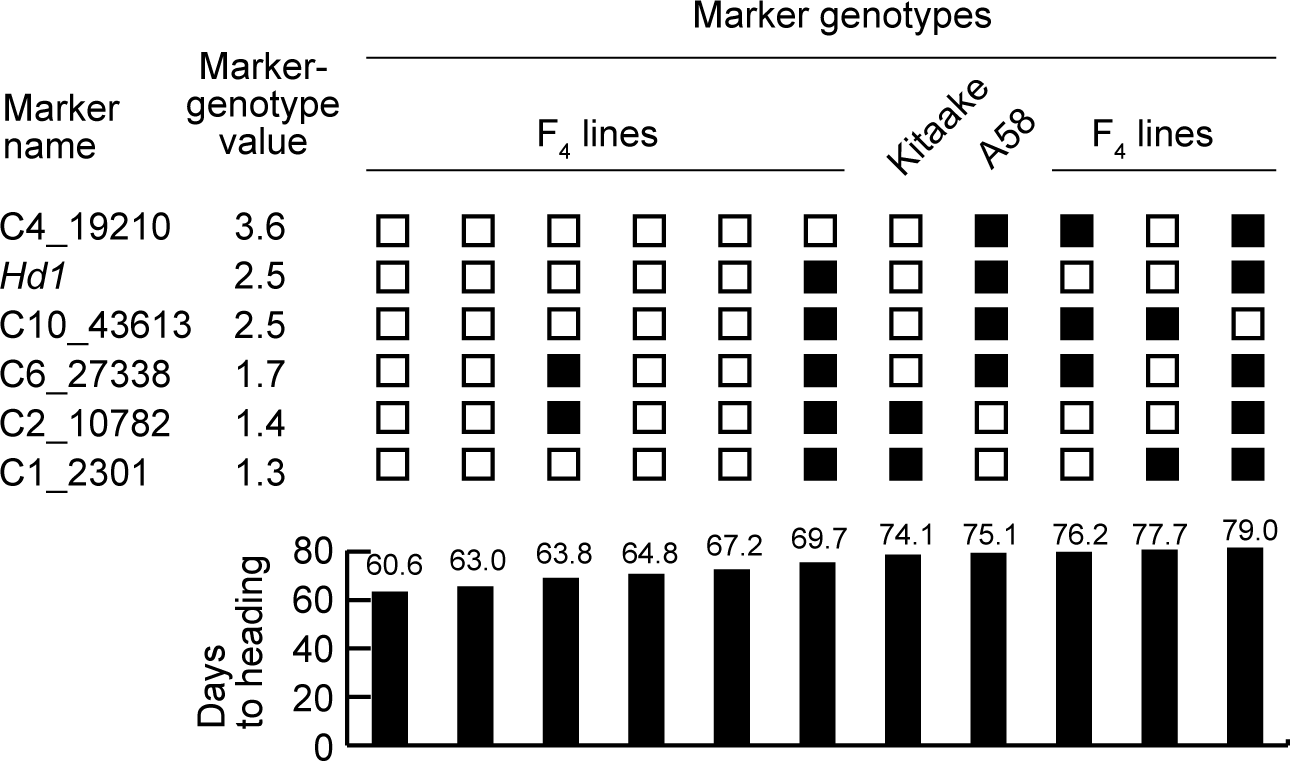
Phenotypic relationships with combinations of six marker genotype values. Among the 30 F4 lines, nine retained the homozygous alleles in the six loci that corresponded to the SNP clusters with QTLs for DTH and Hd1. The effect of each locus on DTH was weighted according to marker genotype values (see Materials and Methods) based on DTH in 2016. Larger values indicate a stronger effect on DTH. Empty squares indicate shorter DTH effects relative to black squares. Kitaake contained four shorter DTH alleles in Ch 4, Hd1, Ch 10, and Ch 6, whereas A58 possessed two shorter DTH alleles in Ch 2 and Ch 1. The two parental cultivars, Kitaake and A58, had DTH of 74.1 and 75.1 days, respectively. DTH in the selected F4 lines ranged from 60.6 to 79.0 days. Each marker name indicates the central SNP in the cluster.

## DISCUSSION

Here, we showed that transgressive segregation occurred in the hybrid progenies of two rice varieties, A58 and Kitaake, both of which have short DTH as an adaptation to high-latitude region. Phenotypic variation beyond the parental range was observed in this segregating population and facilitated uncovering of the genetic basis of transgressive segregation and extreme DTH phenotypes. The two parental varieties shared the same genotypes for three known major QTLs *(E1/Hd4/Ghd7, Hd2/OsPRR37*, and *Hd5/DTH8)* (Figure 3), but different alleles for *Se1/Hd1* and several unknown minor QTLs. Such different genotypes in minor QTLs produced new genetic combinations that resulted in transgressive phenotypes of the progenies. QTLs direct either positive or negative actions based on the effect of parental alleles. If negative QTL alleles in either parent are replaced by the positive alleles of the other parent, the progeny could obtain the desired phenotype because of the presence of more positive alleles (de Vicente and Tanksley 1993; Rieseberg *et al.* 1999). Our results appeared to demonstrate this scenario, because we observed allelic complementation at QTLs, and indicate the importance of such “hidden” genetic variations despite close genetic relationships (Hagiwara *et al.* 2006).

We employed SNP analysis with deep sequencing to obtain a sufficient number of markers for the closely related varieties. This was a powerful approach that detected more than 600 genome-wide SNPs between both Hokkaido-adapted varieties (Figure 4). In addition, such similar genetic backgrounds of the two varieties, A58 and Kitaake, facilitated identification of the minor QTLs that shape transgressive early heading by genome-wide SNP analysis.

Our analysis detected five SNP clusters that corresponded to QTLs and the *Hd1* locus, which contributed to DTH differences in the A58 × Kitaake progenies (Table 1 and 2). These QTLs were involved in both the additive and epistatic effects on extreme heading phenotypes (Table 2 and Figure 5). Among the SNP clusters, the strongest effect was explained by the Ch 4 cluster, in which the Kitaake-derived allele(s) caused decreased DTH (Table 2). The Ch 4 cluster was located from 29.8 to 32.4 Mb on Ch 4 (Figure 4), where only one gene, *Rice FLO-LFY homolog (RFL)* (Kyozuka *et al.* 1998), is functionally characterized as a flowering-related gene by the QTL Annotation Rice

Online (Q-TARO) database (http://qtaro.abr.affrc.go.jp/). Similarly, the Ch 6 cluster (from 24.5 to 25.5 Mb) included a gene for photoperiod sensitivity, *Se5* (Izawa *et al.* 2000). However, no functional polymorphisms in A58 and Kitaake were detected in either *RFL* or *Se5.* In the other QTLs found in the SNP clusters on Chs 1, 2, and 10, no known DTH-related genes were identified. These results demonstrated that some unknown genes present in these SNP clusters affected DTH of the Hokkaido varieties. Interestingly, our analysis also showed possible epistatic interactions between genes in SNP clusters on Chs 1 and 10 that shortened DTH (Figure 5). It was previously thought that epistasis is unlikely to be a major cause of transgressive phenotypes (de Vicente and Tanksley 1993; Rieseberg *et al.* 1999); however, in our study, epistatic interactions explained the transgressive phenotypes observed in the segregating populations (Figure 5).

To date, four genes *(E1/Hd4/Ghd7, Hd2/OsPRR37, Se1/Hd1*, and *Hd5/DTH8)* were reported to control DTH in improved rice varieties in Hokkaido (Ichitani *et al.* 1997; Fujino and Sekiguchi 2005a; Fujino and Sekiguchi 2005b; Nonoue *et al.* 2008; Fujino *et al.* 2013; Koo *et al.* 2013). Among the four genes, loss of functional alleles in *E1/Hd4/Ghd7* and *Hd2/OsPRR37* are necessary to obtain photoperiod insensitivity in rice varieties in northern areas (Fujino and Sekiguchi 2005a; Fujino and Sekiguchi 2005b; Xue *et al.* 2008; Koo *et al.* 2013). Alternatively, the other two genes *(Se1/Hd1* and *Hd5/DTH8)* have small effects on photoperiod insensitivity among the varieties in Hokkaido (Ichitani *et al.* 1997; Fujino and Sekiguchi 2005b; Nonoue *et al.* 2008). In this study, *Se1/Hd1* sequences revealed differences between A58 and Kitaake; A58 has insertion/deletion mutations, whereas Kitaake has the functional allele (Figure 3). Both varieties had the same loss-of-function alleles in the *E1/Hd4/Ghd7* and *Hd2/OsPRR37* loci and had the same functional allele in the *Hd5/DTH8* locus (Figure 3). These results indicated that the improved varieties in Hokkaido might have inherited the same *E1/Hd4/Ghd7, Hd2/OsPRR37*, and *Hd5/DTH8* alleles from a landrace similar to A58, which facilitated adaptation, because photoperiod insensitivity was essential for adaptation. In addition to these four loci, newly identified minor QTLs were identified for DTH on Chs 1, 2, 4, 6 and 10 (Table 2 and Figure 4). These QTLs likely contribute to extreme phenotypes of short and long DTH produced by transgressive segregation based on the composition of their complementary alleles (Figure 6).

Rieseberg *et al.* (1999) made several predictions regarding the cause of transgressive segregation, one of which was consistent with our results: transgressive segregation would likely be observed in the F2 population of parents with more proximal phenotypes (Figure 7). Among the known alleles at the four known loci that are necessary for DTH adaptation to Hokkaido, A58 and Kitaake shared the same alleles at three loci, but not *Hd1*, which indicates that these two varieties possess a considerable amount of common alleles that shorten DTH. Our results demonstrate that transgressive segregation mainly occurred as a result of a few unknown QTLs, in which alleles combined in a complementary manner (Figure 7).

**Figure 7.**
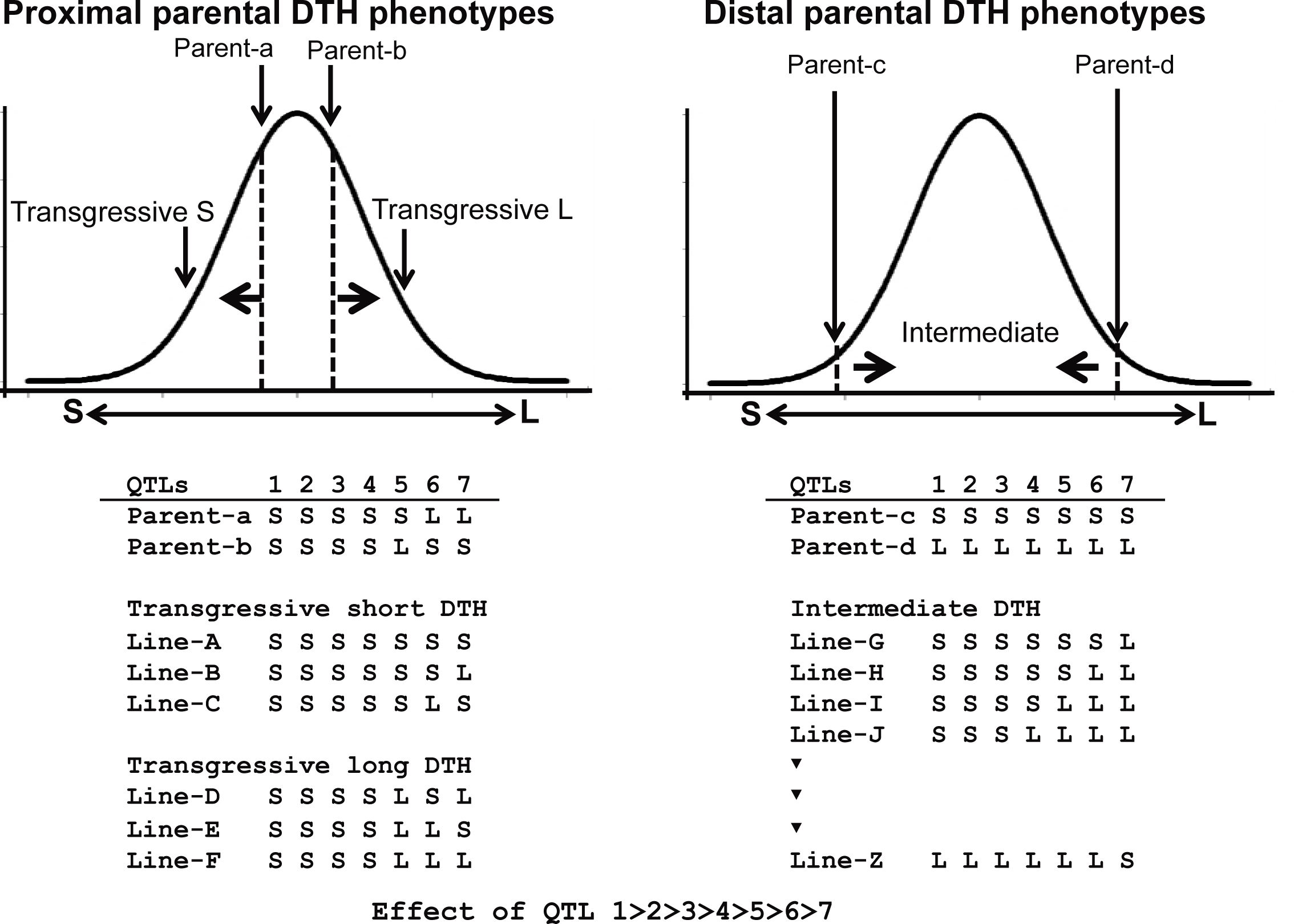
Model of different segregation patterns that occurred in the F2 populations derived from two parental combinations of proximal and distal DTH phenotypes. The left panel represents the segregation pattern of the F2 population between parent-a and -b with proximal DTH phenotypes due to the similar genotypes with a few differences. Because of differences in a few alleles with minor effects on DTH, the F2 progenies produced transgressive phenotypes. The right panel represents the F2 population produced by parents with distal phenotypes and opposite genotypes shows intermediate segregation between both parents. Most of the F2 progenies with mixed genotypes of the parental alleles did not have DTH phenotypes that exceeded those of the parental phenotypes. There are seven loci involved in DTH, and their effects on DTH are ordered as 1 >>> 7. S and L indicate the effect of an allele at each locus that makes DTH shorter or longer, respectively.

According to Rieseberg *et al.* (1999), transgressive segregation tends to occur more frequently in intraspecific crosses, inbred populations, and domesticated populations compared with interspecific crosses, outbred populations, and wild populations, respectively. The lack of a strong positive correlation was observed between parental genetic divergence and transgression frequency (Rieseberg *et al.* 1999; Rieseberg *et al.* 2003). Our previous study (Ota *et al.* 2014) showed that the hybrid progenies of two varieties with distal DTH adapted to different environments exhibited few instances of transgressive DTH phenotypes (Figure 1). Because the parents were adapted to different environments with different genetic backgrounds (e.g., interspecies), a number of the new allelic combinations were generated in the hybrid progenies (Figure 7). Such complex allelic combinations might generate positive and negative genetic interactions, and offset allelic effects. Stochastically, if there is a large number of segregating loci, individuals rarely accumulate only the alleles with positive effects, but most usually contain alleles with negative effects (Figure 7).

This study showed that a few genes and their combinations expanded variation of the DTH phenotype despite similar genetic backgrounds. Consequently, it might be useful to identify QTLs or allelic interactions associated with the transgressive DTH phenotypes in progenies of other varieties with proximal phenotypes. Similarly, to integrate other transgressive phenotypes into breeding programs, alleles with additive effects of minor QTLs should be targeted in varieties with proximal phenotypes.

## ACKNOWLEDGMENTS

We are grateful to Ms. K. Aoyama (Lab. Plant Breeding, Hokkaido University) for her help conducting this study. Kitaake seeds were kindly supplied by the Agricultural Research Department of Hokkaido Research Organization. This work was supported by Takano Life Science Research Foundation and JSPS KAKENHI Grant Number 23658001 (to Y. Kishima), and the Program for Fostering Researchers for the Next

Generation of the Consortium Office for Fostering of Researchers in Future Generations (COFRe), Hokkaido University (to Y. Koide). We thank Mallory Eckstut, PhD, from Edanz Group (www.edanzediting.com/ac) for editing a draft of this manuscript.

## FIGURE LEGENDS

**Figure S1** Experimental scheme of this study. ddRAD-Seq was carried out to detect differences in allele frequency of genome-wide SNPs between early- and late-heading F4 lines.

**Figure S2** Frequency distribution of DTH in the F3 population derived from the A58 × Kitaake cross.

**Figure S3** Effect of the Ch 4 SNP cluster on DTH in 2015 and 2016. Average DTH values of F4 lines with homozygous A58- and Kitaake-type alleles of the SNP cluster on Ch 4, which is represented by the central SNP in the cluster, C4_19396. Significant differences (P < 0.01) between average DTH values were observed in both 2015 and 2016.

**Figure S4** Genetic interactions between SNP clusters on Chs 1, 4, 6, and 10 relative to DTH in 2015. Average DTH values (y-axis) of F4 lines with each genotype are shown in squares. Six combinations of any two loci among the SNP clusters on Chs 1, 4, 6 and 10 are depicted A58-type alleles (black line) and Kitaake-type alleles (gray line) are paired with another locus corresponding to A58-type alleles (the left side) or Kitaake-type alleles (the right side) in x-axis, respectively. An allelic interaction (non-additive interaction) was detectable where the black line crossed the gray line.

## Literature Cited

Alonso-Blanco, C., and B. Mendez-Vigo, 2014 Genetic architecture of naturally occurring quantitative traits in plants: an updated synthesis. Current Opinion in Plant Biology 18: 37–43.

Baird, N. A., P. D. Etter, T. S. Atwood, M. C. Currey, A. L. Shiver et al., 2008 Rapid SNP Discovery and Genetic Mapping Using Sequenced RAD Markers. Plos One 3.

Bolger, A. M., M. Lohse and B. Usadel, 2014 Trimmomatic: a flexible trimmer for Illumina sequence data. Bioinformatics 30: 2114–2120.

Brambilla, V., J. Gomez-Ariza, M. Cerise and F. Fornara, 2017 The Importance of Being on Time: Regulatory Networks Controlling Photoperiodic Flowering in Cereals. Frontiers in Plant Science 8.

Catchen, J. M., A. Amores, P. Hohenlohe, W. Cresko and J. H. Postlethwait, 2011 Stacks: Building and Genotyping Loci De Novo From Short-Read Sequences. G3-Genes Genomes Genetics 1: 171–182.

de Vicente, M. C., and S. D. Tanksley, 1993 Qtl Analysis of Transgressive Segregation in an Interspecific Tomato Cross. Genetics 134: 585–596.

Dittrich-Reed, D. R., and B. M. Fitzpatrick, 2013 Transgressive Hybrids as Hopeful Monsters. Evol Biol 40: 310–315.

Ebana, K., T. Shibaya, J. Z. Wu, K. Matsubara, H. Kanamori et al., 2011 Uncovering of major genetic factors generating naturally occurring variation in heading date among Asian rice cultivars. Theoretical and Applied Genetics 122: 1199–1210.

Fujino, K., and H. Sekiguchi, 2005a Mapping of QTLs conferring extremely early heading in rice (Oryza sativa L.). Theoretical and Applied Genetics 111: 393–398.

Fujino, K., and H. Sekiguchi, 2005b Identification of QTLs conferring genetic variation for heading date among rice varieties at the northern-limit of rice cultivation. Breeding Science 55: 141–146.

Fujino, K., U. Yamanouchi and M. Yano, 2013 Roles of the Hd5 gene controlling heading date for adaptation to the northern limits of rice cultivation. Theoretical and Applied Genetics 126: 611–618.

Goddard, M. E., and B. J. Hayes, 2007 Genomic selection. Journal of Animal Breeding and Genetics 124: 323–330.

Hagiwara, W. E., K. Onishi, I. Takamure and Y. Sano, 2006 Transgressive segregation due to linked QTLs for grain characteristics of rice. Euphytica 150: 27–35.

Harlan, J. R., 1976 Genetic Resources in Wild Relatives of Crops. Crop Science 16: 329–333.

Hori, K., K. Matsubara and M. Yano, 2016 Genetic control of flowering time in riceintegration of Mendelian genetics and genomics. Theoretical and Applied Genetics 129: 2241–2252.

Huang, X. H., and B. Han, 2014 Natural Variations and Genome-Wide Association Studies in Crop Plants. Annual Review of Plant Biology, Vol 65 65: 531–551.

Ichitani, K., Y. Okumoto and T. Tanisaka, 1997 Photoperiod sensitivity gene of Se-1 locus found in photoperiod insensitive rice cultivars of the northern limit region of rice cultivation. Breeding Science 47: 145–152.

Ishiguro, S., K. Ogasawara, K. Fujino, Y. Sato and Y. Kishima, 2014 Low Temperature-Responsive Changes in the Anther Transcriptome’s Repeat Sequences Are Indicative of Stress Sensitivity and Pollen Sterility in Rice Strains. Plant Physiology 164: 671–682.

Izawa, T., T. Oikawa, S. Tokutomi, K. Okuno and K. Shimamoto, 2000 Phytochromes confer the photoperiodic control of flowering in rice (a short-day plant). Plant Journal 22: 391–399.

Koo, B. H., S. C. Yoo, J. W. Park, C. T. Kwon, B. D. Lee et al., 2013 Natural Variation in OsPRR37 Regulates Heading Date and Contributes to Rice Cultivation at a Wide Range of Latitudes. Molecular Plant 6: 1877–1888.

Kyozuka, J., S. Konishi, K. Nemoto, T. Izawa and K. Shimamoto, 1998 Down-regulation of RFL, the FLO/LFY homolog of rice, accompanied with panicle branch initiation. Proceedings of the National Academy of Sciences of the United States of America 95: 1979–1982.

Langmead, B., and S. L. Salzberg, 2012 Fast gapped-read alignment with Bowtie 2. Nature Methods 9: 357–U354.

Nonoue, Y., K. Fujino, Y. Hirayama, U. Yamanouchi, S. Y. Lin et al., 2008 Detection of quantitative trait loci controlling extremely early heading in rice. Theoretical and Applied Genetics 116: 715–722.

Ota, Y., S. Ishiguro, E. Aoyama, R. Aiba, R. Iwashiro et al., 2014 Isolation of a major genetic interaction associated with an extreme phenotype using assorted F2 populations in rice. Molecular Breeding 33: 997–1003.

Peterson, B. K., J. N. Weber, E. H. Kay, H. S. Fisher and H. E. Hoekstra, 2012 Double Digest RADseq: An Inexpensive Method for De Novo SNP Discovery and Genotyping in Model and Non-Model Species. Plos One 7.

Rick, C. M., and P. G. Smith, 1953 Novel Variation in Tomato Species Hybrids. American Naturalist 87: 359–373.

Rieseberg, L. H., M. A. Archer and R. K. Wayne, 1999 Transgressive segregation, adaptation and speciation. Heredity 83: 363–372.

Rieseberg, L. H., A. Widmer, A. M. Arntz and J. M. Burke, 2002 Directional selection is the primary cause of phenotypic diversification. Proceedings of the National Academy of Sciences of the United States of America 99: 12242–12245.

Rieseberg, L. H., A. Widmer, A. M. Arntz and J. M. Burke, 2003 The genetic architecture necessary for transgressive segregation is common in both natural and domesticated populations. Philosophical Transactions of the Royal Society of London Series B-Biological Sciences 358: 1141–1147.

Takata, M., Y. Kishima and Y. Sano, 2005 DNA methylation polymorphisms in rice and wild rice strains: Detection of epigenetic markers. Breeding Science 55: 57–63.

Tanksley, S. D., and S. R. McCouch, 1997 Seed banks and molecular maps: Unlocking genetic potential from the wild. Science 277: 1063–1066.

Thompson, J. D., D. G. Higgins and T. J. Gibson, 1994 Clustal-W - Improving the Sensitivity of Progressive Multiple Sequence Alignment through Sequence Weighting, Position-Specific Gap Penalties and Weight Matrix Choice. Nucleic Acids Research 22: 4673–4680.

Vega, U., and K. J. Frey, 1980 Transgressive Segregation in Inter and Intraspecific Crosses of Barley. Euphytica 29: 585–594.

Xue, W. Y., Y. Z. Xing, X. Y. Weng, Y. Zhao, W. J. Tang et al., 2008 Natural variation in Ghd7 is an important regulator of heading date and yield potential in rice. Nature Genetics 40: 761–767.

Yano, M., Y. Katayose, M. Ashikari, U. Yamanouchi, L. Monna et al., 2000 Hd1, a major photoperiod sensitivity quantitative trait locus in rice, is closely related to the arabidopsis flowering time gene CONSTANS. Plant Cell 12: 2473–2483.

